# S51 family peptidases provide resistance to peptidyl-nucleotide antibiotic McC

**DOI:** 10.1101/2022.03.27.485901

**Authors:** Eldar Yagmurov, Konstantin Gilep, Marina Serebryakova, Yuri I. Wolf, Svetlana Dubiley, Konstantin Severinov

## Abstract

Microcin C-like compounds are natural Trojan horse peptide-nucleotide antibiotics produced by diverse bacteria. The ribosomally-synthesized peptide parts of these antibiotics are responsible for their facilitated transport into susceptible cells. Once inside the cell, the peptide part is degraded, releasing the toxic payload, an isoaspartyl-nucleotide that inhibits aspartyl-tRNA synthetase, an enzyme essential for protein synthesis. Bacteria that produce microcin C-like compounds have evolved multiple ways to avoid self-intoxication. Here, we describe a new strategy through the action of S51 family peptidases, which we name MccG. MccG cleaves the toxic isoaspartyl-nucleotide rendering it inactive. While some MccG homologs are encoded in gene clusters responsible for McC-like compounds biosynthesis, most are encoded by stand-alone genes whose products may provide basal level of resistance to peptide-nucleotide antibiotics in phylogenetically distant bacteria.

**SIGNIFICANCE:** We identified a natural substrate for a major phylogenetic clade of poorly characterized S51 family proteases from bacteria. We show that these proteins can contribute to basal level of resistance to an important class of natural antibiotics.

## INTRODUCTION

*E. coli* peptidyl-nucleotide antibiotic microcin C (McC) is a prototypical compound of a distinct class of ribosomally synthesized and post-translationally modified peptides (RiPP) (1). McC-like compounds biosynthesis is encoded in *mcc* biosynthetic gene clusters (BGCs) found in the genomes of numerous gram-negative and gram-positive bacteria (2, 3). The minimal set of genes required for production of an McC-like compound comprises *mccA,* which encodes a precursor peptide, *mccB,* coding for the ThiF-like nucleotidyltransferase, and a gene whose product is responsible for antibiotic export. Other genes frequently present in *mcc* BGCs are responsible for either additional decorations on the nucleotide part of the final product or for self-immunity of the producer. Upon translation, the MccA peptide is modified by MccB with the formation of peptidyl-nucleotide in which the C-terminal asparagine is converted to isoasparagine and linked to a nucleoside monophosphate through a non-hydrolyzable phosphoramide linkage (4) (Figure 1A). The reaction consumes two molecules of triphosphates per one molecule of McC synthesized and proceeds through a stable peptidyl-succinimide intermediate nucleoside (5). While the peptide parts of different McCs vary greatly in their sequence and length (though the C-terminal asparagine is strictly conserved), their nucleotide parts so far are found in only two distinct forms, adenosine or cytosine monophosphates with or without additional decorations (4, 6) (Figure 1B).

**Figure 1.**
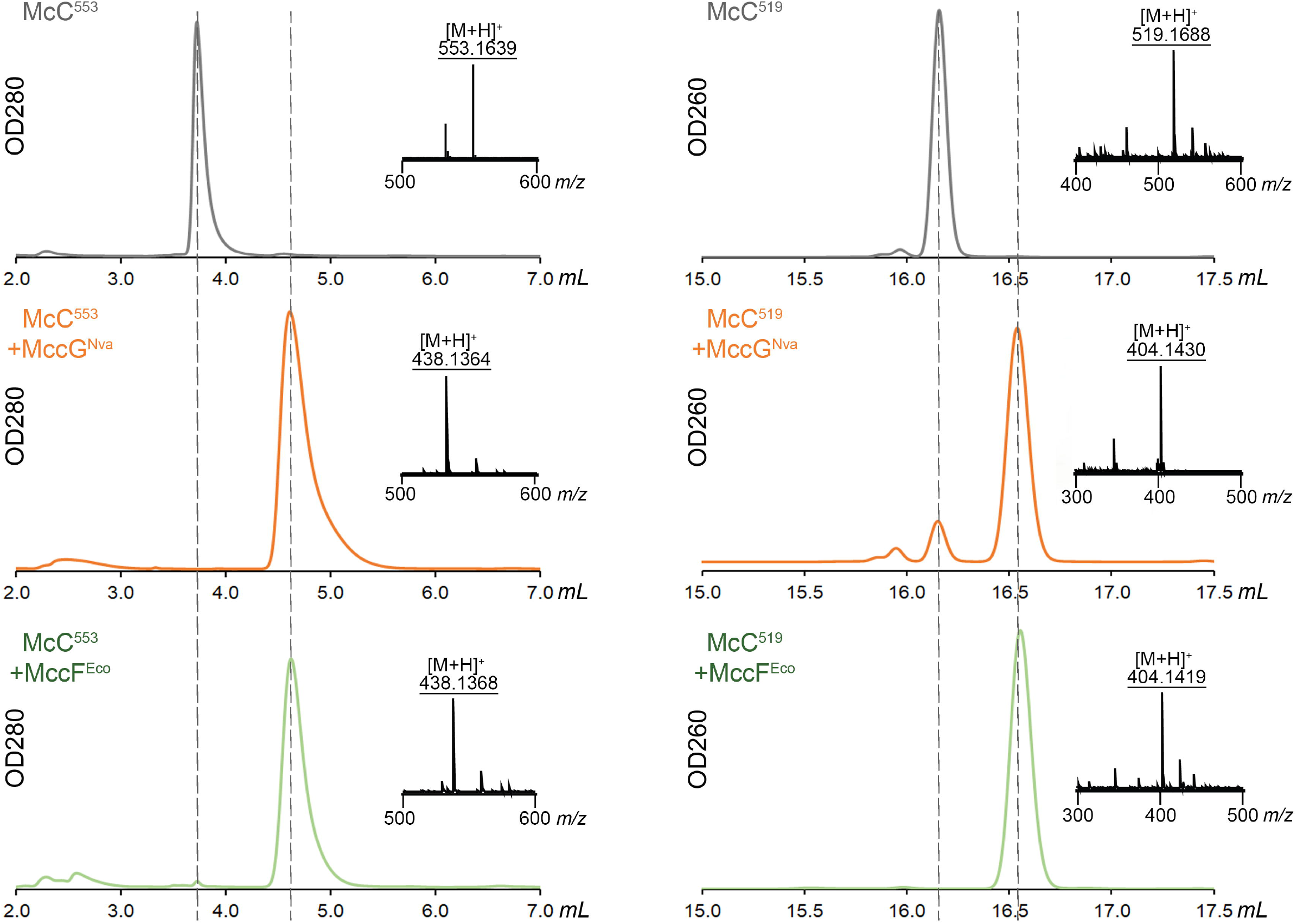
Microcin C biosynthesis, mode of action, and structural diversity of McC-like compounds. **(A)** McC^Eco^ biosynthesis by the producing cell containing the *mcc* operon; its uptake by a susceptible cell, and the inhibitory action of the compound on AspRS are schematically shown. The *mccA* gene codes for a precursor peptide, *mccB* encodes a nucleotidyltransferase, *mccC* - an export MFS transporter, *mccD* and *mccE* are required for aminopropyl group installation and self-immunity (*mccE*). **(B)** The structure of McC^Eco^, a peptidyl-adenylate, and peptidyl-cytidylate McC^Yps^ from *Yersinia pseudotuberculosis*. The nitrogen atom participating in the formation of non-hydrolyzable phosphoramide linkage is shown in red; the aminopropyl modification is shown in blue. The position of the carboxymethyl group on the cytidyl moiety of McC^Yps^ is currently unknown. The intermediate of the AspRS-catalyzed reaction of tRNAA^sp^ charging (aspartyl-AMP) is shown on the right. McC^519^ and McC^553^ correspond to processed forms of McC^Eco^ and McC^Yps^, respectively.

The *E. coli* McC (McC^Eco^) inhibits the growth of *E. coli* and closely related bacteria lacking the *mcc* operon using a Trojan-horse mechanism (7). The compound is actively imported into the target cell through oligopeptide transporter YejABEF (8). In the cytoplasm, the seven aminoacid-long peptide part is proteolytically processed by aminopeptidases PepA, PepB, and PepN with the release of the active payload – isoasparaginyl-adenosine monophosphate decorated with an aminopropyl group at the phosphate (9). This “processed McC^Eco^” is a structural mimic of aspartyl-adenosyl monophosphate, an intermediate of the tRNA^Asp^ charging reaction catalyzed by aspartyl-tRNA synthetase (AspRS). Binding of processed McC in the active center of AspRS results in protein biosynthesis inhibition (7), stringent response (10), and, eventually, cell death. All McC-like compounds studied to date have the Trojan-horse mechanism of action and target AspRS (7, 11). The conservation of essential genes in *mcc* clusters and the fact that all MccA precursor peptides contain a terminal asparagine residue imply that both the mechanism and the intracellular target is conserved for all compounds of this class. Indeed, cytosine containing McC-like compounds also target the AspRS (6, 11).

While mature McC-like compounds are exported from the producing cell by dedicated transporters, a certain amount is inevitably processed releasing the toxic payload inside the producer (10). Several strategies to prevent self-intoxication of producers have been described. The Gcn5-related N-acetyltransferase MccE2 acetylates the primary amino group of processed McC^Eco^ thus preventing its binding to AspRS (12). L,D-carboxypeptidase MccF hydrolyses the carboxamide bond between the peptide and nucleotide parts of intact McC^Eco^ or aminoacyl and nucleotide parts of processed McC (13). Phosphoramidase MccH from *Hyalangium minutum* hydrolyzes the phosphoramide linkage of processed McC (14).

In this study, we report a new protein from *Nocardia vaccinii mcc*-like gene cluster that provides immunity to McC-like compounds. We name this protein MccG. MccG belongs to a superfamily of S51 peptidases. The type member of this family is peptidase E (PepE) – an aspartyl dipeptidase that hydrolyzes the peptide bond in dipeptides with an N-terminal L-aspartate (15). We show that MccG from *Nocardia vaccinii* and its homologs hydrolyze the carboxamide bond in processed forms of McC-like compounds with the formation of aspartate and a modified nucleotide phosphoramide. We further show that MccG homologs frequently encoded as stand-alone genes in various bacterial genomes can protect cells from McC-like compounds. Our findings reveal a new strategy of resistance to toxic aminoacyl-nucleotides and underscore the diversity of strategies that have been harnessed in the course of evolution to address the problem of self-intoxication of producers of McC-like RIPPs.

## MATERIALS AND METHODS

### DNA manipulation, molecular cloning and protein purification

All cloning steps were conducted in *E. coli* DH5*α* (F^−^ φ80lacZΔM15 Δ(lacZYA-argF) U169 recA1 endA1 hsdR17(rK^−^, mK^+^) phoA supE44 λ–thi-1 gyrA96 relA1). Protein purification was performed in *E. coli* BL21DE3 (F^−^ompT hsdSB (rB^−^, mB^−^) gal dcm (DE3)). Phusion High-fidelity DNA polymerase (Thermo Scientific, USA) was used for DNA amplification. DNA primer synthesis and DNA sequencing were performed by Evrogen (Russia). For the list of primers used in the study refer to Table S1. Unless indicated otherwise, the genomic DNA was used as a template for PCR amplification.

For production of recombinant MccB^Nva^ enzyme, PCR amplified *mccB^Nva^* gene was digested with BamHI and SacI restriction endonucleases and introduced into pRSFDuet-1 vector digested with the same endonucleases. The resultant pRSF_6xHis_*mccB^Nva^* contained a sequence encoding MccB^Nva^ enzyme fused with an N-terminal hexahistidine affinity tag.

For *in vivo* spot toxicity assay, pBAD_SalRBS (14) vector was digested by SalI and HindIII and combined with the PCR amplified fragments of the corresponding genes digested with same restriction endonucleases. Genes encoding MccG-homologs from *Rothia dentocariosa* ATCC17931, *Mycobacteroides abscessus* subsp. *massiliense* strain 616, and *Vibrio parahaemolyticus* ATCC17802 were purchased from IDT, USA as synthetic DNA fragments. To create mutants of MccG^Nva^, site-directed mutagenesis was employed using overlap extension PCR (16).

For recombinant protein purification of MccF^Eco^, PepE^Eco^, MccG^Nva^ its homologs and mutants, the PCR amplified genes encoding corresponding enzymes and lacking stop codon were digested with NdeI and XhoI and introduced into a linearized with same restriction endonucleases pET22 vector (Novagen-Millipore, USA) with an engineered sequence encoding a C-terminal hexahistidine affinity tag.

For protein production *E. coli* BL21DE3 strain transformed with an appropriate plasmid was grown in 250 mL of TB medium supplemented with kanamycin at 37 °C, and moderate constant shaking until OD_600_ ~ 0.6. Upon reaching OD_600_ ~ 0.6, the bacterial culture was induced with 0.5 mM IPTG and grown at 18 °C for additional 16 h. The cells were harvested by centrifugation, resuspended in buffer A [20 mM Tris-HCl (pH 8.0), 150 mM NaCl, 2 mM imidazole, 5% glycerol], supplemented with 0.5 mM PMSF and disrupted by sonication. The resultant lysate was cleared by centrifugation at 30 000 x *g* for 20 min and applied to Talon CellThru Co^2+^ chelating resin (TaKaRa-Clontech). The protein bound resin was washed with 3 column volumes (CV) of buffer A, and 5 CV of buffer B [20 mM Tris-HCl (pH 8.0), 50 mM NaCl, 20 mM imidazole]. The protein was eluted with 2 CV of buffer C [20 mM Tris-HCl (pH 8.0), 50 mM NaCl, 300 mM imidazole, 10% glycerol]. The fractions containing the eluted protein were analyzed by SDS-PAGE and bands corresponding to purified proteins were subjected to tryptic digestion with subsequent MALDI-TOF mass spectrometry identification of the protein. Following elution, the protein was dialysed in buffer D [20 mM Tris-HCl (pH 8.0), 50 mM NaCl, 10% glycerol], and stored at −80 °C until further use.

### *In vitro* nucleotidylation and carboxymethylation reactions

To recreate the McC^Nva^ biosynthesis *in vitro,* the nucleotidyl transfer and carboxymethylation reactions were conducted using the synthetic MccA^Nva^ peptide (MKIVLKLKRIVRGAGPIIVSN), chemically synthesized cxSAM (6, 17) and a recombinant MccB^Nva^ protein. The reaction containing 100 μM of chemically synthesized MccA^Nva^ peptide, 2 mM equimolar mix of nucleotide triphosphates, 2 mM cxSAM, and 5 μM recombinant MccB^Nva^ in a reaction buffer [50 mM Tris-HCl, pH 8.0, 150 mM NaCl, 5 mM MgCl_2_, 5 mM DTT] was incubated for 16 hours at 28 °C, then stopped by addition of TFA to a concentration 0.1% (v/v). The products of the reaction were analyzed by MALDI-TOF mass spectrometry for the presence of peptidyl-carboxymethylcytidine.

### *In vivo* McC toxicity assay

*E. coli* strain B transformed with pBAD_SalRBS vector containing *mccF^Eco^, pepE^Eco^, mccG^Nva^* or its homologs. Transformants were selected on the LB plates containing was 100 μg/mL ampicillin. A single colony was inoculated into 2xYT medium supplemented with 100 μg/mL ampicillin and 10 mM arabinose and grow at 37 °C for 16 h. The overnight culture was diluted 1000-fold in M9 agar medium, supplemented with 1% glycerol, 0.1% yeast extract, 10 mM arabinose and 100 μg/mL ampicillin. 3 μL drops of McC^Eco^ and McC^Yps^ of varying concentration were placed on the surface of the agar plate and allowed to dry. Plates were incubated for 16 h at 30 °C to form a lawn.

### Preparation of McC^553^ and McC^519^

Purification of McC^Eco^ and McC^Yps^ was performed as described previously (11). HPLC purified McC^Yps^ and McC^Eco^ were dissolved in 3 mL of the reaction buffer [50 mM HEPES, pH 7.4, 0.75 mM MnCl_2_, and 5 mM TCEP] to a concentration of 100 μM. The solution was then combined with recombinant PepA, PepB and PepN from *E. coli* to a concentration of 10 μM each and the mixture was incubated at 32 °C for 16 h. Upon incubation, the reaction was quenched by the addition of an equal volume of 100% acetonitrile. The pH of the solution was adjusted to 2.0 by addition of TFA and the solution was incubated for 30 min on ice to allow the proteins to precipitate. After incubation, the protein precipitate was removed by centrifugation at 16 000 x *g* for 15 min (4 °C) and the supernatant was collected and lyophilized. The protein pellet was then extracted 2 times with a 50% acetonitrile solution in deionized water and the extracts were also lyophilized.

The supernatant and extract fractions were dissolved in 0.1% TFA solution in deionized water and applied to Agilent Prep 5 C18 column (10 x 250 mm, particle size 5 μm, Agilent Technologies). The processed McC^Yps^ and McC^Eco^ were first purified in 0.1% TFA/acetonitrile system in a linear 1-13% gradient of acetonitrile, followed by a second round of purification in ammonium acetate buffer, pH 4.3 in linear 1-13% gradient of acetonitrile on YMC-Triart C18 column (150 x 3 mm, particle size 3 *μ*m, YMC).

### Chemical synthesis of Asp-pNA

Asp-pNA was synthesized from protected Fmoc-Asp(Ot-Bu)-OH (Sigma) by coupling with 4-niroaniline (pNA) (Sigma) as described in Nedev *et al.,* 1993 (18) with minor modifications. Briefly, Fmoc-Asp(Ot-Bu)-OH was activated with isobutyl chloroformate in tetrahydrofuran (THF), which facilitated subsequent reaction with weakly nucleophilic 4-nitroaniline. Fmoc deprotection was performed with Tris(2-aminoethyl)amine (TAEA) as described in Peterson et al., 2010 (19). Side chain carboxyl group was deprotected from t-Bu in TFA:CH_2_Cl_2_ (19:1) for 30 min. Solvent was removed by rotary evaporation. Solid product was washed with ethyl either, lyophilized, resuspended in 0.1% TFA in MQ and applied to Agilent 5 Prep-C18 RP-HPLC column (250 x 10 mm, particle size 5 *μ*M, Agilent Technologies). The purification of Asp-pNA was carried out in a linear 5-25% gradient of acetonitrile; fractions absorbing at 313 nm were analyzed for the presence of Asp-pNA by ESI mass spectrometry. The fractions containing a validated Asp-pNA were subjected to the additional chromatographic purification on the same columns in a linear 15-20% gradient of acetonitrile in 30 mM ammonium acetate buffer, pH 4.3.

### *In vitro* hydrolysis assays

To test the hydrolytic activity of MccF^Eco^, PepE^Eco^, MccG^Nva^ its homologs and mutants, 1 *μ*M of the respective recombinant enzyme was combined with 100 micromolar McC^Yps^, McC^Eco^ and their processed forms (McC^553^ and McC^519^, respectively) in the reaction buffer [50 mM HEPES (pH 7.4), 2 mM MgCl_2_, 100 mM NaCl, 2 mM DTT]. The mixture was incubated for 30 minutes at 28 °C then the reaction was quenched by addition of TFA to a concentration of 0.1% (v/v). The reaction products were analyzed by RP-HPLC followed by ESI-MS analysis.

To test peptidase E activity of enzymes, 200 *μ*M of Asp-pNA were ed with 2 *μ*M of the recombinant enzyme in the reaction buffer [50 mM HEPES (pH 7.4), 2 mM MgCl_2_, 100 mM NaCl, 2 mM DTT]. The hydrolysis was carried out for 30 minutes at 28 °C then stopped by addition of TFA. The hydrolysis of Asp-pNA was monitored by the increase in optical density at OD_410_ corresponding to the accumulation of released pNA, using the Nanodrop 2000c spectrophotometer (Thermo Scientific, USA).

### RP-HPLC analysis of the products of *in vitro* reactions

The analysis of enzymatic reactions was performed on Infinity II 1260 Liquid Chromatography system (Agilent). Separation of the enzymatic reaction products occurred on YMC-Triart C18 column in 30 mM ammonium acetate buffer system (pH 4.3) in a linear gradient of acetonitrile from (0-10%) for 25 minutes. Processing of chromatograms was performed in OpenLab CDS ChemStation, Agilent. The elution profiles were exported in comma separated value (.csv) format for visualization.

### Mass spectrometry analysis

High resolution electrospray ionization mass spectrometry (ESI-MS) analysis. Q-TOF Maxis Impact II (Bruker Daltonics) mass-spectrometer with electrospray ionization was used for sample analysis. Lyophilized HPLC fractions from biochemical reactions were dissolved in 0.1% formic acid in deionized water and introduced directly to the instrument using the syringe pump. The spectra were recorded in a positive ion mode in a range from 100 *m/z* to 750 *m/z*. The temperature of ion source was 200 °C, pressure of carrier gas 2.5 bar, the gas flow 5 ml/min, the voltage at the capillary 4 kV. The fragmentation spectra were recorded in AutoMS mode.

Matrix Assisted Laser Desorption Time of Flight (MALDI-TOF) mass spectrometry. Sample aliquots were combined with the matrix mix (Sigma-Aldrich) on a steel target. The mass spectra were recorded on an UltrafleXtreme MALDI-TOF/TOF mass spectrometer (Bruker Daltonics) equipped with a neodymium laser. The molecular MH+ ions were measured in reflector mode; the accuracy of the measured results was within 0.1 Da.

### Phylogenetic analysis of S51 family peptidases

Sequence alignments of S51 family peptidases from the NCBI CDD database (cd03129, cd03145, and cd03146) were used as queries in PSI-BLAST (20) searches against a database, containing 21.4 M protein sequences encoded in 13,116 completely sequenced archaeal and bacterial genomes, available as of March 2019 (obtained from the NCBI FTP site, <https://ftp.ncbi.nlm.nih.gov/genomes/ASSEMBLY_REPORTS/>). Sequences, matching these profiles, were clustered using MMSEQS2 (21) with the similarity threshold of 0.5; sequences within each cluster were aligned using MUSCLE (22). Alignments were compared to each other using HHSEARCH (23) an iteratively merged using HHALIGN, guided by an UPGMA tree, constructed from the matrix of HHSEARCH scores. The approximate maximum likelihood tree was reconstructed using FastTree (24) with WAG evolutionary model and gamma-distributed site rates.

## RESULTS

During bioinformatical screening of sequenced bacterial genomes we identified a group of *mcc-* like biosynthetic gene clusters (BGCs) in *Nocardia vaccinii* NBRC 15922 and various *Mycobacteroides abscessus* strains (2). Despite some differences in their architecture, these clusters closely resemble the previously characterized *mcc* operon from *Yersinia pseudotuberculosis* IP32953 (11) (Figure 2A). We decided to focus our study on *N. vaccinii mcc*.

**Figure 2.**
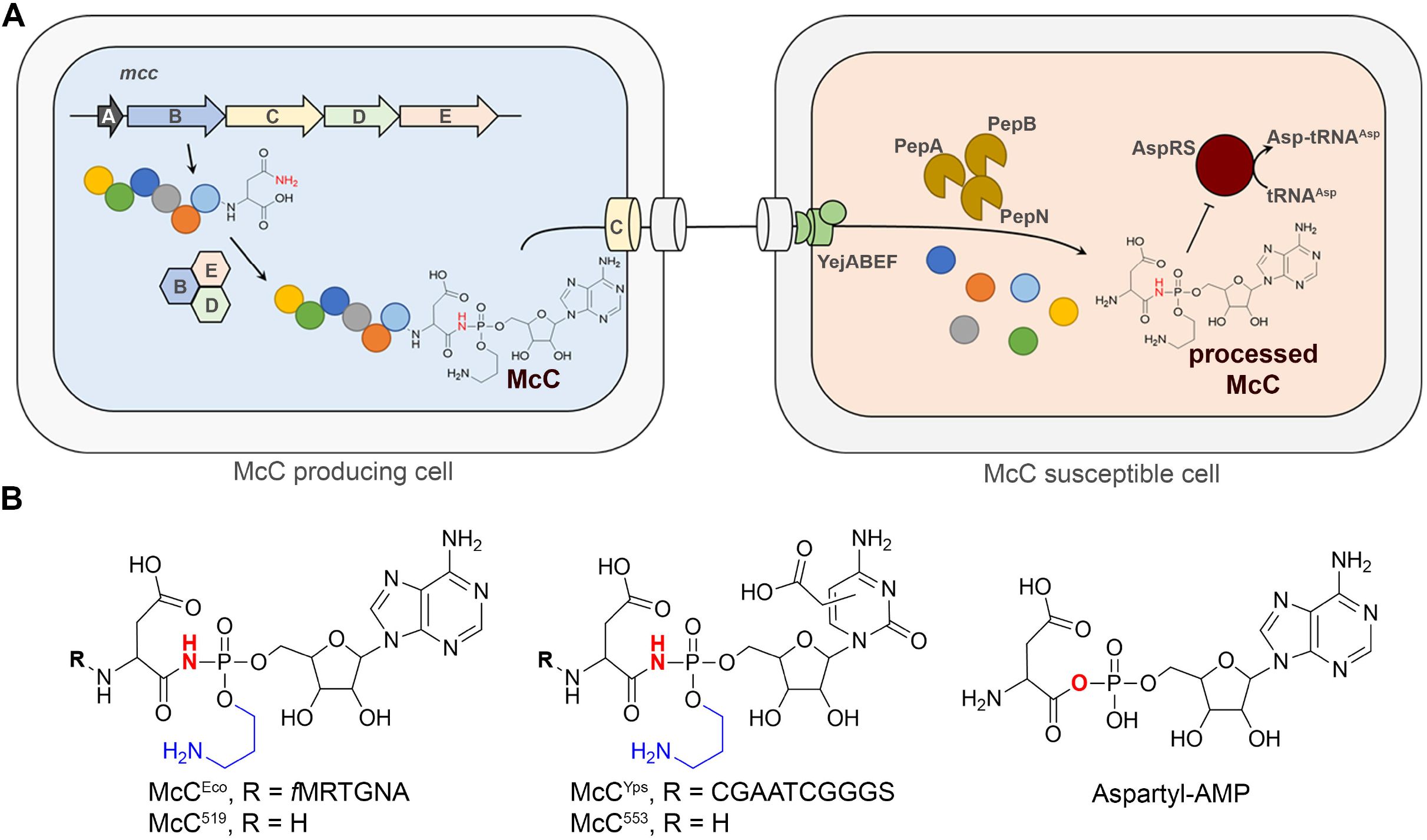
The *mcc*-like cluster from *N. vaccinii* NBRC 15922 (locus tags NV1_RS35535-NV1_RS35560) and its products. **(A)** Organization of the *mcc*-like BGCs from *E. coli, Y. pseudotuberculosis* IP 32953, *N. vaccinii* NBRC 15922, and *M. abscessus* subsp*. massiliense*. The general functions of the genes are indicated by colors and discussed in the text. The arrows with the numbers in between aligned BGCs indicate the degree of identity of amino acid sequences of *N. vaccinii* and *M. abscessus mcc* gene products. **(B)** The proposed mechanism of posttranslational modifications of MccA^Nva^. The carboxymethyl and the (carboxy)aminopropyl groups are shown in green and blue, respectively, and the nitrogen atom involved in the phosphoramide bond is shown in red.

The *mcc* cluster from *N. vaccinii* contains a putative *mccA* gene, which codes for a 21 amino acid long precursor peptide, and *mccB^Nva^*, which codes for a protein similar to *Yersinia pseudotuberculosis* MccB (MccB^Yps^). Unlike the *E. coli* nucleotidyltransferase MccB, MccB^Yps^ and MccB^Nva^ are binfunctional proteins containing an N-terminal nucleotidyltransferase domain and a C-terminal carboxymethyl transferase domain. The product of *mccS^Nva^* is homologous to MccS^Yps^, a carboxy-S-adenosylmethionine (cxSAM) synthase (11). The products of *mccD^Nva^* and *mccE_1_^Nva^* are homologous to enzymes that jointly install the aminopropyl decoration at the phosphate group of peptidyl nucleotides in *E. coli* and *Y. pseudotuberculosis* (11, 25). A homolog of *mccX^Yps^*, which encodes a protein of unknown function, is also present in the *N. vaccinii mcc* operon. The *N. vaccinii mcc* cluster contains an additional gene, *mccG^Nva^*, located between the *mccC* and *mccB* genes. Such a gene is absent from the *Y. pseudotuberculosis mcc* operon (Figure 2A), but is present in the identical location of the *M. abscessus* clusters.

By analogy with *Y. pseudotuberculosis* microcin C (McC^Yps^) one can assume that *N. vaccinii* McC (McC^Nva^) comprises the MccA peptide modified with carboxymethylated cytidylate additionally decorated with an aminopropyl moiety (Figure 2B). To test this hypothesis, we partially reconstructed the proposed pathway *in vitro* using the synthetic MccA^Nva^ precursor and the recombinant MccB^Nva^ (Figure S1). MALDI-MS analysis of the reaction products allowed identification of MccA^Nva^-cytidylate and MccA^Nva^-carboxymethylated cytidylate, thus confirming that MccA^Nva^ is cytidylated and carboxymethylated by MccB^Nva^.

The *Y. pseudotuberculosis mcc* operon encodes MccE2^Yps^, a self-immunity enzyme that acetylates the amino group of processed McC, making it unable to inhibit AspRS. No such enzyme is encoded by the *N. vaccinii mcc* operon. In fact, no genes coding for any known immunity proteins are present in *N. vaccinii mcc*. We therefore hypothesized that the product of *mccG* gene, a gene that is absent in previously characterized *mcc* operons, can perform the immunity function in the *N. vaccinii mcc* BGC. MccG^Nva^ belongs to the MEROPS S51 family of peptidases (26), which includes PepE form *S. enterica* (27) and CphB cyanophycinase from *Synechocystis* sp. PCC6803 (28) that hydrolyze Asp-Xaa dipeptides and multi-*L*-arginyl-poly-(*L*-aspartic acid) cyanophycin, respectively. We hypothesized that the product of *mccG^Nva^* gene encodes a peptidase that cleaves peptidyl-nucleotides.

To test the proposed immunity function, we cloned *mccG^Nva^* into an arabinose-inducible pBAD vector and transformed the resulting plasmid pBAD-*mccG^Nva^* into McC-susceptible *E. coli* cells. Cells transformed with pBAD-*mccF^Eco^* plasmid expressing the self-immunity L,D-carboxypeptidase MccF from *E. coli mcc* BGC were used as a positive control. Cells harboring an empty pBAD vector served as a negative control.

To check if *E. coli* cells expressing *mccG^Nva^* acquired resistance to peptidyl-nucleotides, drops of solution of McC^Eco^ and McC^Yps^ were deposited on the surface of freshly seeded cell lawns and formation of growth inhibition zones was monitored after overnight incubation of plates at 30 ^o^C [see Materials and Methods]. As can be seen from Figure 3, both McC^Yps^ and McC^Eco^ inhibited the growth of *E. coli* cells harboring the empty vector. In contrast, expression of *mccF^Eco^* provided resistance to both McCs, as expected. Cells containing pBAD-*mccG^Nva^* were fully resistant to McC^Yps^, and partially resistant to McC^Eco^. We conclude that MccG^Nva^ is indeed a self-immunity enzyme capable of inactivating modified peptidyl-nucleotides, with a higher apparent preference towards peptidyl-cytidylates.

**Figure 3.**
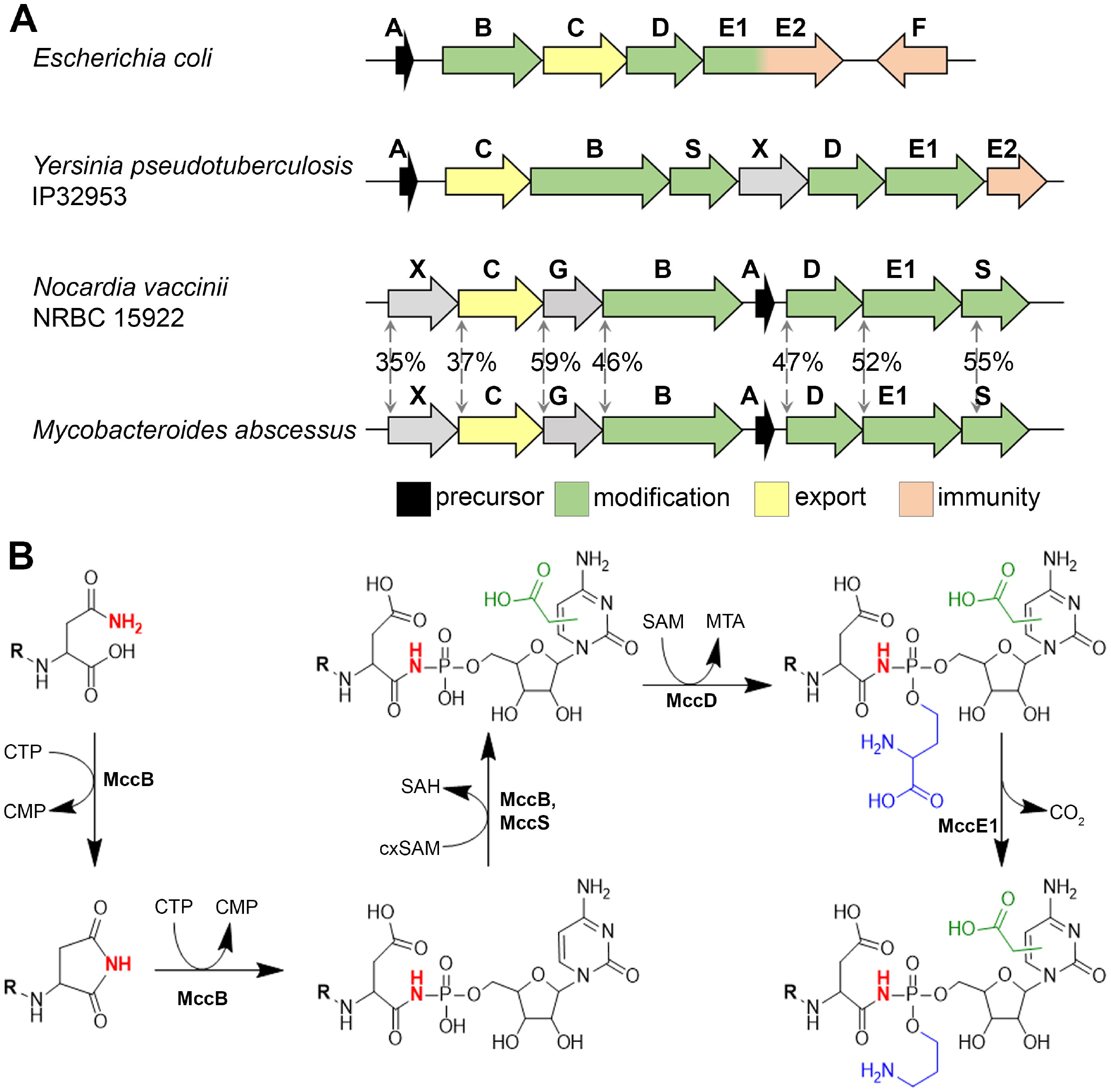
Overexpression of *mccG^Nva^* renders *E. coli* cells resistant to peptidyl-nucleotides. Drops of McC^Yps^ and McC^Eco^ solutions of indicated concentrations were deposited on the surface of freshly seeded lawns of McC-susceptible *E. coli* cells transformed with indicated plasmids. Photographs of plates after overnight growth at 30 ^o^C at conditions of induction of cloned plasmid-borne genes are shown.

To identify the mechanism of immunity provided by MccG^Nva^, the recombinant MccG^Nva^ and MccF^Eco^ proteins were incubated with McC^Yps^ and McC^Eco^. The reaction products were separated using RP-HPLC and the content of chromatographic peaks was analyzed with high resolution ESI-MS. No changes to the original compounds were observed when McC^Yps^ and McC^Eco^ were incubated with MccG^Nva^ (Figure S2). In contrast, MccF^Eco^ efficiently cleaved off the peptide parts of both McC^Yps^ and McC^Eco^ as confirmed by accumulation of the [M+H]^+^ mass ions at *m/z* 438.14 and 404.14, respectively (Figure S2). We therefore conclude that mature McC^Yps^ and McC^Eco^ are not substrates for MccG^Nva^.

To infer the mechanism of protective activity of MccG^Nva^, a phylogenetic tree of S51 family peptidases, using 3,097 protein sequences found in 13,116 completely sequenced archaeal and bacterial genomes, was constructed. The resulting tree contains six major clades (Figure 4). The previously studied PepE aspartyl dipeptidase from *Salmonella enterica* (27) and cyanophycinase form *Synechocystis* sp. PCC6803 (28) belong to clades I and IV, respectively (Figure 4A). Several proteins that do not possess peptidase activity and have no established physiological function belong to clade II (Figure 4A) (29). MccG^Nva^, together with PepE^Bs^ from *Bacillus subtilis* (29) belongs to clade V. PepE^Bs^ was previously shown to hydrolyze the model substrate of aspartyl dipeptidases, L-aspartic acid α-(p-nitroanilide) (Asp-pNA), *in vitro* (29), however, its natural substrates are unknown.

**Figure 4.**
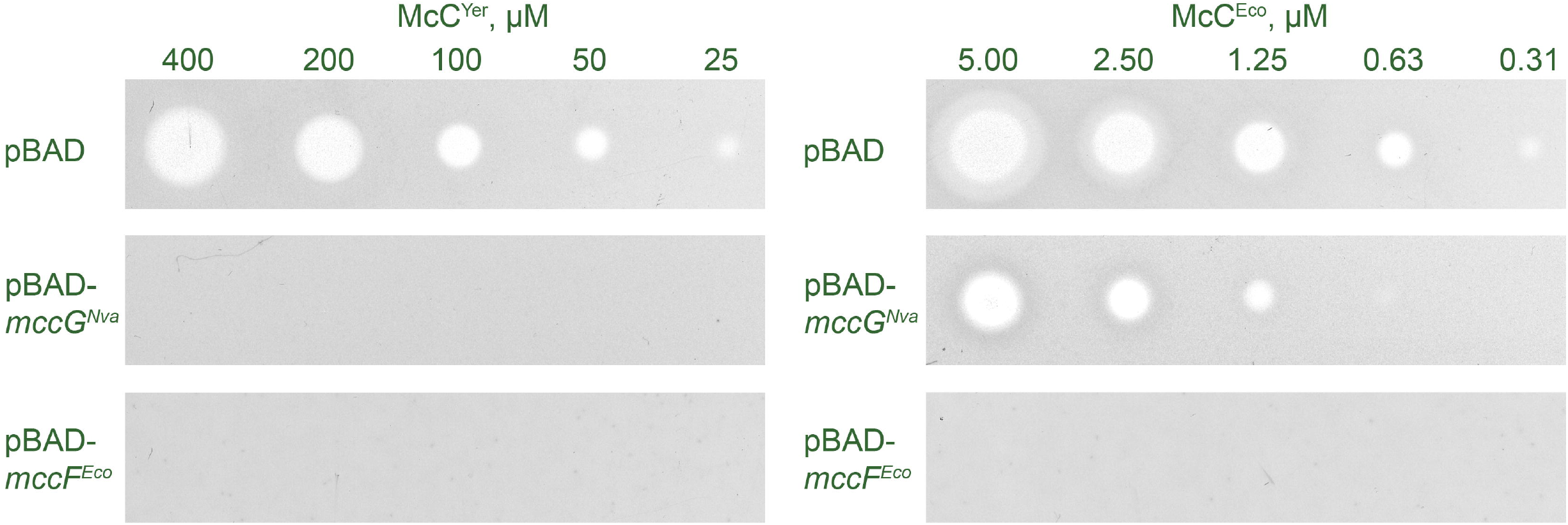
Phylogenetic analysis of MccG and its homologs from different bacteria. **(A)** A phylogenetic tree of aspartyl dipeptidases. The previously characterized representatives of S51 family peptidases (15, 28, 29) are numbered from 1 through 6 and highlighted with open circles. The *N. vaccinii* MccG (marked by a green circle and number 10) is located on a distinct clade, marked as V. Several MccG homologs from clade V are numbered from 1 through 9 and highlighted with green circles. **(B)** Resistance of McC-susceptible *E. coli* cells overproducing indicated plasmid-borne MccG homologs to McC^Yps^ and McC^Eco^. Resistance was calculated as a ratio of minimal inhibitory concentration (MIC) value for McC^Yps^ and McC^Eco^ obtained on lawns of *E. coli* cells transformed with plasmids expressing an MccG homolog to the MIC value obtained on lawns of cells transformed with an empty pBAD vector.

To check if the ability to confer resistance to McC-like compounds is unique to MccG^Nva^, we tested MccG^Mab^ from the *M. abscessus mcc* BGC and several MccG^Nva^ homologs from clade V encoded by stand-alone genes. The latter included MccG-like proteins from *Artrobacter* sp. FB24*, Bacillus cohnii* DSM6307*, Bacillus cereus* ATCC4342*, Bacillus subtilis* 168*, Bacillus valismortis* DSM11031*, Bacillus velezensis* FZB42*, Rothia dentocariosa* ATCC17931, and *Vibrio parahamoelyticus* ATCC4342 (Figure 4A). The genes encoding MccG^Nva^ homologs were cloned into the pBAD vector, the resulting plasmids were transformed in McC-susceptible *E. coli* and cells expressing *mccG* homologs were tested for susceptibility to McC^Eco^ and McC^Yps^. Cells overexpressing cloned *E. coli pepE* were also tested.

As can be seen from (Figure 4B, Figure S3), overexpression of *E. coli* dipeptidase PepE did not protect cells from the McC action. To confirm that recombinant PepE is an active aspartyl dipeptidase we have synthesized its substrate Asp-*p*NA (29). Incubation of Asp-*p*NA with recombinant PepE led to hydrolysis of the compound at the carboxamide bond and formation of fluorogenic *p*-nitroaniline (Figure S4). Thus, recombinant PepE^Eco^ is active but does not confer resistance to McC-like antibiotics.

As expected, MccG^Mab^, which is encoded in an *mcc* gene cluster, conferred immunity to both McC^Yps^ and McC^Eco^. Interestingly, five out of seven tested MccG^Nva^ homologs that are encoded by stand-alone genes also provided resistance to McC^Yps^ and McC^Eco^. Expression of *M. abscessus* MccG^Mab^*, B. subtilis* 168 MccG^Bsu^, and *B. velezensis* MccG^Bve^ provided higher levels of resistance to McC^Eco^ than did MccG^Nva^. It remains to be determined whether this result is due to broader specificity of these enzymes or is a trivial effect of higher levels of production in a heterologous system. Two proteins belonging to the basal branches of clade V, MccG^Bce^ and MccG^Vpa^ did not confer immunity to McCs, suggesting that these enzymes have different substrate specificities from the rest of MccG enzymes tested.

To show that the peptidase activity is required for the McC-protecting function of MccG^Nva^, we attempted to predict its aminoacid residues important for catalysis. However, low sequence similarity between the *S. enterica* PepE, *Synechocystis* sp. CphB, and MccG^Nva^ precluded unambiguous identification of the catalytic triad (Figure S5). We therefore performed structure modeling of MccG^Nva^ using AlfaFold v 2.1.0 (30). The monomeric structure was modeled with high confidence (pLDDT >90) for most of the protein sequence except for 20 amino acids at the C-terminus. As expected, the MccG^Nva^ showed the highest structural similarity to cyanophycinase from *Synechocystis* sp. PCC6803 (28) (PDB:3EN0, RMSD 2.6) and PepE from *S. enterica* (31) (PDB:6A4R, RMSD 2.6). The superposition of the of *S. enterica* PepE and MccG^Nva^ monomers revealed an identical architecture of the catalytic center in two enzymes (Figure 5A). The catalytic triad of PepE is composed of Ser120, His157, and Glu192 residues, which form a network of hydrogen bonds that position the nucleophilic Ser to interact with the substrate. However, while the catalytic Ser133 and His168 residues are conserved, the third amino acid in the MccG^Nva^ triad is an aspartate (Asp196). This residue is conserved in clade V enzymes.

**Figure 5.**
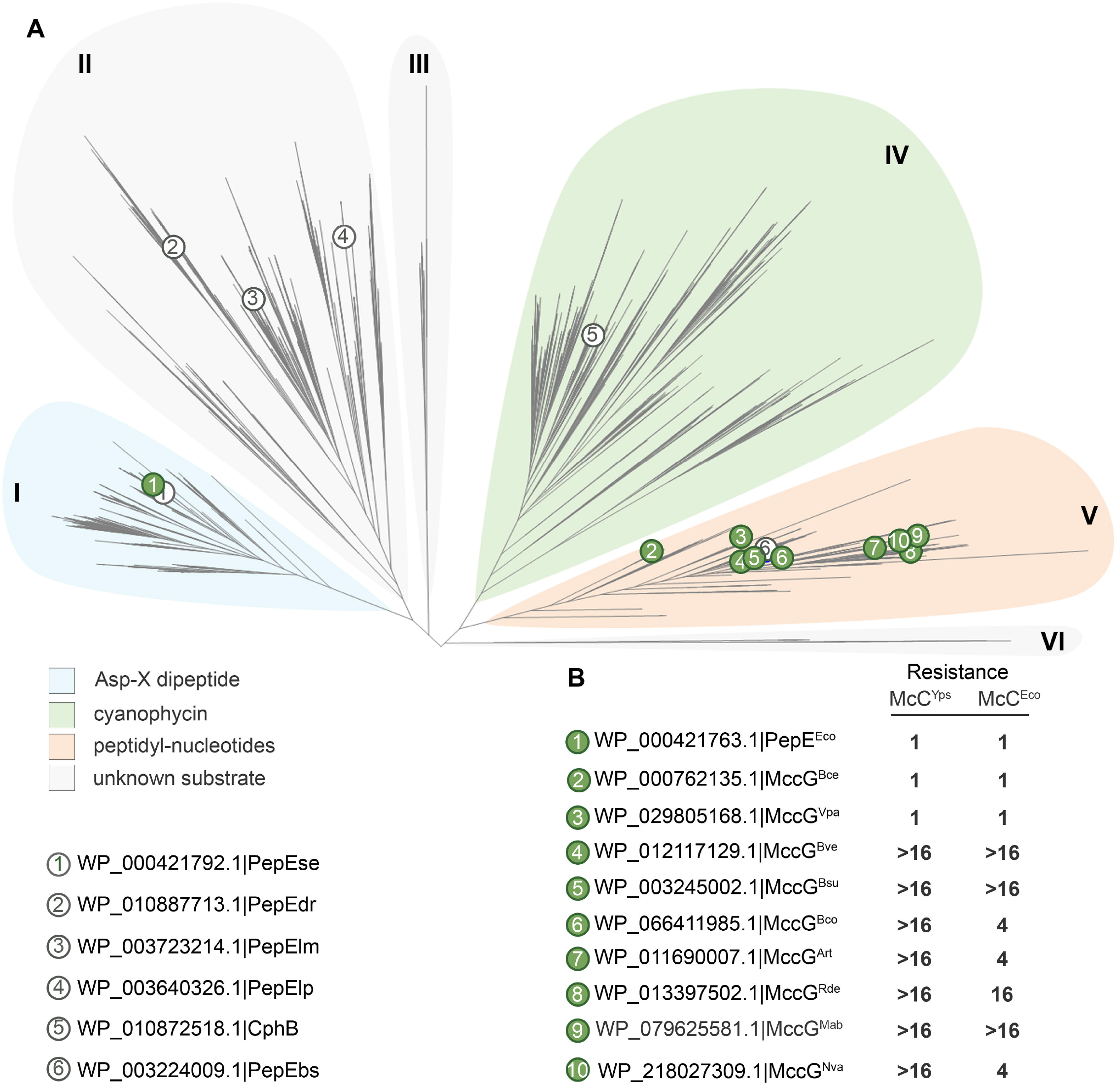
Mutational analysis of MccG^Nva^. **(A)** Superposition of the catalytic centers of PepE aspartyl dipeptidase (PDB:6A4R, shown in grey) and the MccG^Nva^ structural model (shown in green). The catalytic residues are shown in sticks. **(B)** Resistance to McC^Yps^ and McC^Eco^ of *E. coli* cells overproducing indicated MccG^Nva^ mutants.

We used site-specific mutagenesis to substitute each of the conserved amino acid of MccG^Nva^ catalytic triad (Ser133, His168, and Asp196) for alanine. We have also reconstructed the “canonical PepE catalytic triad” in MccG^Nva^, by constructing a D196E single substitution mutant with an expectation that this may affect its ability to cleave processed McCs, which PepE lacks. *E. coli* cells expressing plasmid-borne mutant *mccG^Nva^* were tested for susceptibility to McC^Eco^ and McC^Yps^. Expectedly, Ser133Ala and His168Ala substitutions abolished MccG-mediated protection from McC^Yps^. Expression of MccG^Nva^ with the D196A substitution led to substantially decreased resistance to peptidyl-nucleotide antibiotics, while the D196E mutant was resistant to Mcc^Yps^ (Figure 5B). Compared to wild-type MccG^Nva^, the ability of D196E mutant to protect cells from McC^Eco^ was somewhat compromised, as judged by the size of growth inhibition zones. We conclude that the catalytic activity of MccG^Nva^ is required for protecting cells against inhibitory action of peptidyl-nucleotide antibiotics, however, the presence of an aspartate instead of glutamate found in PepE is not the sole determinant of MccG ability to detoxify processed McC.

We next hypothesized that MccG^Nva^ is a peptidase capable of hydrolyzing the peptide bond solely in processed McCs, small molecules with an aspartate residue coupled to a nucleotide. We prepared aminopropylated forms of carboxymethylated aspartamide-cytidylate (McC^553^) and aspartamide-adenylate (McC^519^) by *in vitro* proteolysis of mature McC^Yps^ and McC^Eco^, respectively, and incubated them with recombinant MccG^Nva^ or MccF^Eco^. The reaction products were separated by RP-HPLC and subjected to ESI-MS analysis. Both MccF^Eco^ and MccG^Nva^ fully converted McC^553^ into a new compound with distinct chromatographic mobility. The ESI-MS analysis of the observed peak revealed a [M+H]^+^ ion at *m/z* 438.1364, matching aminopropylated carboxymethylcytidine phosphoramide (calculated monoisotopic mass of the ion is 438.1384 Da) (Figure 6). The MS/MS fragmentation spectra of the compound verified the assignment (Figure S6).

**Figure 6.**
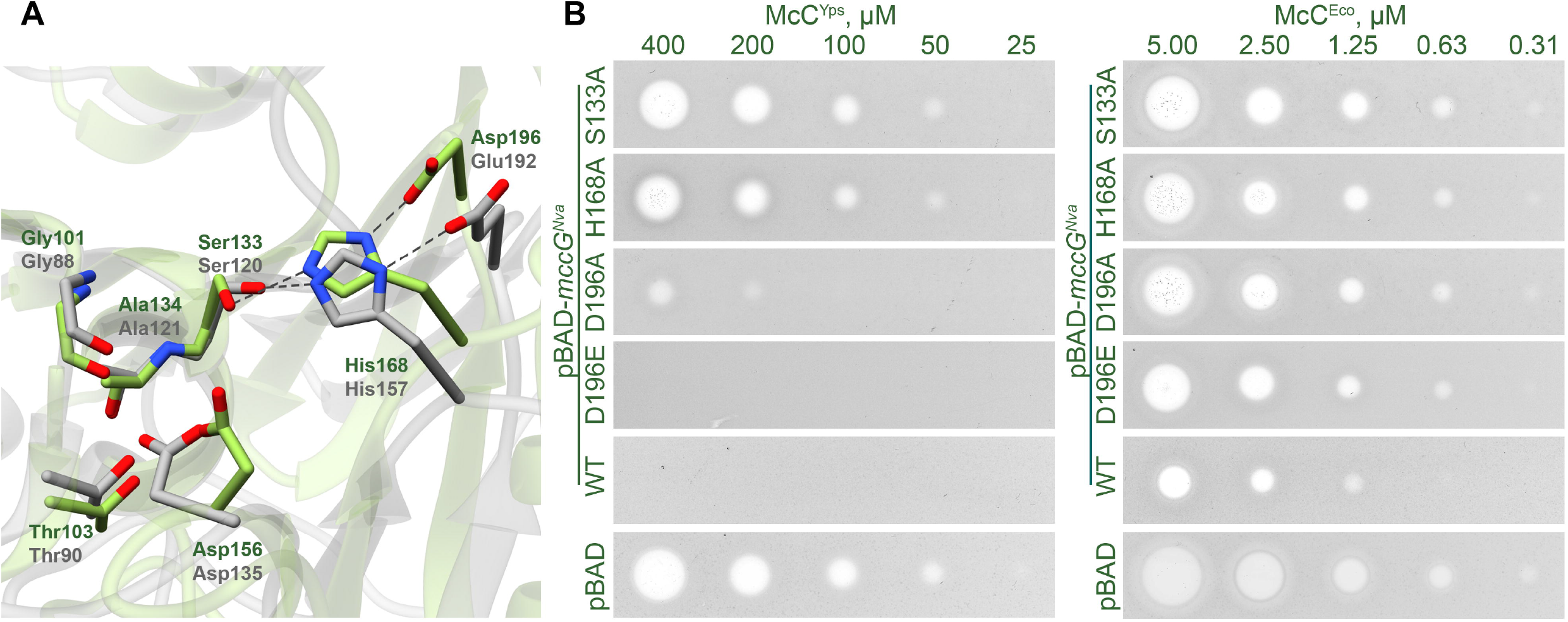
MccG^Nva^ hydrolyzes the carboxamide bond in isoasparginyl-carboxymethylcytidylate McC^553^ and isoasparaginyl-adenylate McC^519^. RP-HPLC elution profiles of the reaction products upon incubation of McC^553^ and McC^519^ without additional enzymes or with MccG^Nva^ or MccF^Eco^. ESI-MS analyses of corresponding chromatographic peaks are superimposed with HPLC elution profiles. [M+H]^+^ at *m/z* 553.1632 corresponds to McC^553^; [M+H]^+^ at *m/z* 519.1688 corresponds to McC^519^; [M+H]^+^ at *m/z* 438.1384 corresponds to aminopropylated carboxymethylcytidine phosphoramide and [M+H]^+^ at *m/z* 404.1449 corresponds to aminopropylated adenosine phosphoramide.

After incubation of McC^519^ with MccG^Nva^ the chromatographic peak corresponding to the initial compound ([M+H]^+^ ion at *m/z* 519.1685) was observed alongside with a ([M+H]^+^ ion at *m/z* 404.1419). The conversion was complete in reactions containing MccF^Eco^. The [M+H]^+^ ion at *m/z* 404.1419 matches aminopropylated adenosine phosphoramide. ESI MS/MS spectrum of the compound confirmed the assignment (Figure S6). Thus, MccG^Nva^ hydrolyzes carboxamide bond in processed McC-like with a clear preference towards aminoacyl cytidylates.

Given the data presented above, we conclude that MccG^Nva^ hydrolyzes the carboxamide bond in processed McC-like compounds through a mechanism that is similar to that used by serine proteases to hydrolyze peptide bonds (Figure S7).

## DISCUSSION

The mechanism of McC-like compounds action necessitates the need for immunity strategies against the compound, whether endogenously produced, or imported from the outside. While minimal *mcc* operons containing just three genes coding for the peptide precursor -- the nucleotidyl transferase that produced the active compound, and the export pump -- are known, many *mcc*-like gene clusters encode enzymes that provide self-immunity to the producing cell (2, 3). Using *in vitro* reconstitution, we here show that the product of *Nocardia vaccinii mcc* is a peptidyl-cytidylate similar to McC^Yps^ (Figure S1) (11). In *Y. pseudotuberculosis mcc*, self-immunity is achieved through the function of MccE2, which covalently modifies processed McC rendering it inactive. In this paper we describe a novel self-immunity enzyme MccG from the *N. vaccinii mcc* operon. We show that MccG is an unusual peptidase that hydrolyzes the carboxamide bond between the aminoacyl and nucleotide moieties of processed McC (Figure 3). MccG homologs encoded by stand-alone genes can also protect cells from McC.

It is not clear at this point which of the two mechanisms of self-immunity is ancestral. While *mccE* genes are found in different locations in *mcc* operons from different sources, the *mccG* genes so far are found in the *N. vaccinii* and *M. abscessus mcc* BGCs and their location is distinct from known locations of *mccE* genes. Be that as it may, comparisons of different BGCs suggest that self-immunity functions can be lost from *mcc* operons and analogous functions can then be regained by recruiting stand-alone genes from the core genome of the bacterial cell. While this has not been explicitly checked, it is possible that minimal three-gene *mcc* operons reside in bacteria which provide the self-immunity functions by stand-alone MccE2-, MccG-, MccH-, and MccF-like enzymes that mop up toxic processing products of *mcc* operons accumulating inside the producers, whether from intracellularly produced compounds or from those imported from the outside. Recruitment of any one of these genes or their combinations provides a clear advantage for mobile genetic elements that often carry *mcc* operons and allow them to efficiently colonize diverse hosts without decreasing their fitness.

Unlike MccF, a previously described McC self-immunity peptidase (12), MccG^Nva^ can only detoxify processed forms of McC-like compounds (Figure 4, Figure S2). While MccF cleaves both modified peptidyl-adenylates and cytidylates efficiently, MccG from *N. vaccinii* has a clear preference for peptidyl-cytidylates (Figure 4). Yet, M. *abscessus* MccG^Mab^, *B. subtilis* 168 MccG^Bsu^, and *B. velezensis* MccG^Bsu^ provided higher levels of resistance to McC^Eco^ than MccG^Nva^. It is thus possible that some MccG homologs are as versatile as MccF and can protect cells from both cytosine and adenosine containing McC-like compounds.

## Acknowledgements

This work was supported by NIH RO1 grant AI117270 (to KS and Satish A. Nair), the Russian Science Foundation grant RSF #19-14-00266 to SD, and a grant from the Ministry of Science and Higher Education of Russian Federation (agreement No. 075-10-2021-114 from 11 October 2021 to KS). Y.I.W. is supported through the intramural program of the U.S. National Institutes of Health. MALDI MS facility became available to us in the framework of the Moscow State University Development Program PNG 5.13. LC-MS experiments were carried out at the Institute of Biomedical Chemistry using equipment of “Human proteome” Core Facility Centre (Moscow, Russia).

## Supplementary materials

**Figure S1**. (A) MALDI MS spectra of synthetic MccA^Nva^ precursor peptide (upper panel) and the products of its *in vitro* modification by recombinant MccB^Nva^ (lower panel). The *in vitro* coupled nucleotidylation-carboxymethylation reaction was performed in the presence of a chemically synthesized cxSAM and equimolar mix of four NTPs (11). [M+H]^+^ at *m/z* 2305.5 corresponds to unmodified MccA^Nva^; mass-ions at *m/z* 2610.5 and at *m/z* 2668.5 match the MccA^Nva^-cytidylate and carboxymethylated MccA^Nva^-cytidylate, respectively. [M+Na]^+^ at *m/z* 2632.5 marked with an asterisk corresponds to a sodium adduct of MccA^Nva^-cytidylate. (B) MALDI TOF MS/MS spectrum of the products of MccA^Nva^ *in vitro* modification by MccB^Nva^ ([M+H]^+^ ions at *m/z* 2668.5). Mass shifts for 44 and 58 Da correspond to removal of carboxyl and carboxymethyl groups, respectively. Mass differences in 111 and 194 Da match the loss of cytosine nucleobase and monophosphorylated ribose, correspondingly. [M+H]^+^ ion at *m/z* 2287 matches dehydrated MccA peptide (MKIVLKLKRIVRGAGPIIVSN).

**Figure S2**. Intact McC-like compounds are not substrates for MccG^Nva^. RP-HPLC elution profiles of McC^Yps^ ([M+2H]^2+^ at *m/z* 658.2061 and [M+3H]^3+^ at *m/z* 439.1395) and McC^Eco^ ([M+2H]^2+^ at *m/z* 589.2312 and [M+3H]^3+^ at *m/z* 393.1566) alone or after incubation with MccG^Nva^ and MccF^Eco^.

**Figure S3**. Susceptibility to McC^Yps^ and McC^Eco^ of *E. coli* harboring the plasmids with the indicated *mccG^Nva^* homologs.

**Figure S4**. Recombinant PepE^Eco^ hydrolyzes synthetic Asp-*p*NA substrate. Absorbance spectra of Asp-pNA (left panel) and Asp-pNA incubated with PepE^Eco^ (right panel).

**Figure S5.** A multiple sequence alignment of cyanophycinase from *Synechocystis* sp. (WP_010872518.1), the PepE peptidase from *S. enterica* (WP_000421792.1), and MccG^Nva^ from *N. vaccinii* NBRC 15922 (WP_218027309.1) is presented. The alignment was built using MUSCLE with default parameters. Experimentally confirmed catalytic residues of Cph and PepE and are highlighted in red. Based on the sequence alignment, only two of these residues, a serine and a histidine are conserved in all three proteins.

**Figure S6.** ESI-MS/MS fragmentation spectra of the products of MccG^Nva^-mediated hydrolysis of the McC^519^ (upper panels) and McC^553^ (lower panels). Parent ions on mass-spectra are labeled with red color font.

**Figure S7. A proposed reaction mechanism of MccG**. At the first step, Ser133 deprotonated with His168 attacks the carbonyl group of processed McC aspartate forming the first tetrahedral complex. The His168 complex with proton is stabilized via a hydrogen bond with the negatively charged Asp196. The His168 hydrogen is then taken by the nitrogen of the phosphamide bond of the substrate, releasing the nucleotide part. Next, the remaining covalently bound aspartate is attacked by a water molecule, which, similarly to Ser133 at the first step, is deprotonated by His168. The newly formed tetrahedral complex decomposes with the release of free aspartate and the catalytic triad is returned to its initial state.

**Table S1**. Primers used in the study.

