## Supplementary materials for "S51 family peptidases provide resistance to peptidyl-nucleotide antibiotic McC"

Konstantin Severinov

### Supplementary materials

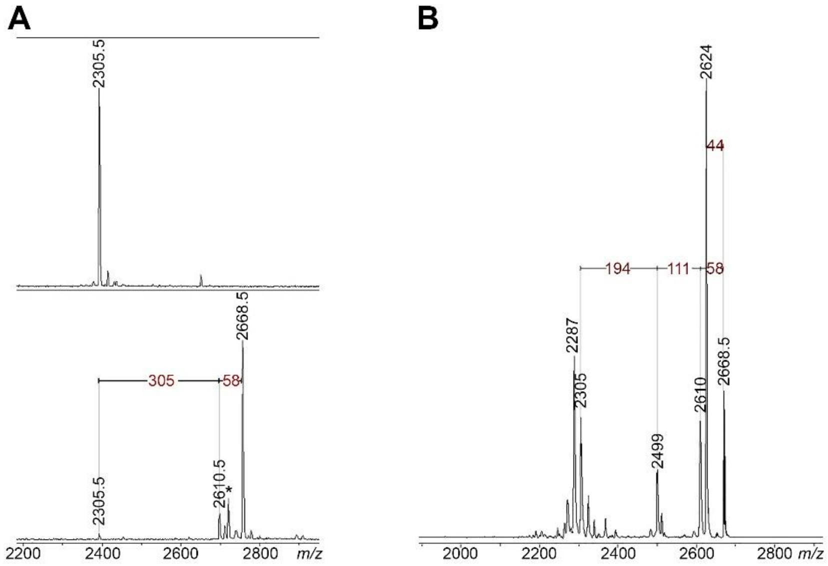


**Figure S1. (A)** MALDI MS spectra of synthetic MccA$^{Nva}$ precursor peptide (upper panel) and the products of its *in vitro* modification by recombinant MccB$^{Nva}$ (lower panel). The *in vitro* coupled nucleotidylation-carboxymethylation reaction was performed in the presence of a chemically synthesized cxSAM and equimolar mix of four NTPs (11). $[M+H]^+$ at *m/z* 2305.5 corresponds to unmodified MccA$^{Nva}$; mass-ions at *m/z* 2610.5 and at *m/z* 2668.5 match the MccA$^{Nva}$-cytidylate and carboxymethylated MccA$^{Nva}$-cytidylate, respectively. $[M+Na]^+$ at *m/z* 2632.5 marked with an asterisk corresponds to a sodium adduct of MccA$^{Nva}$-cytidylate. **(B)** MALDI TOF MS/MS spectrum of the products of MccA$^{Nva}$ *in vitro* modification by MccB$^{Nva}$ ($[M+H]^+$ ions at *m/z* 2668.5). Mass shifts for 44 and 58 Da correspond to removal of carboxyl and carboxymethyl groups, respectively. Mass differences in 111 and 194 Da match the loss of cytosine nucleobase and monophosphorylated ribose, correspondingly. $[M+H]^+$ ion at *m/z* 2287 matches dehydrated MccA peptide (MKIVLKLKRIVRGAGPIIVSN).

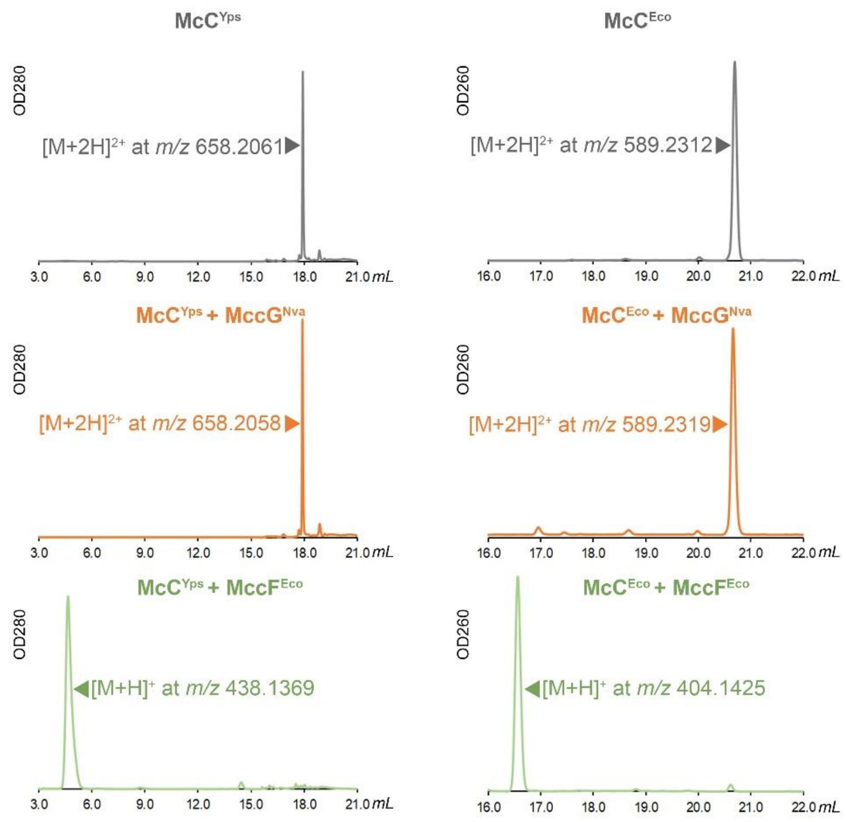


**Figure S2.** Intact McC-like compounds are not substrates for $MccG^{Nva}$. RP-HPLC elution profiles of $McC^{Yps}$ ($[M+2H]^{2+}$ at *m/z* 658.2061 and $[M+3H]^{3+}$ at *m/z* 439.1395) and $McC^{Eco}$ ($[M+2H]^{2+}$ at *m/z* 589.2312 and $[M+3H]^{3+}$ at *m/z* 393.1566) alone or after incubation with $MccG^{Nva}$ and $MccF^{Eco}$.

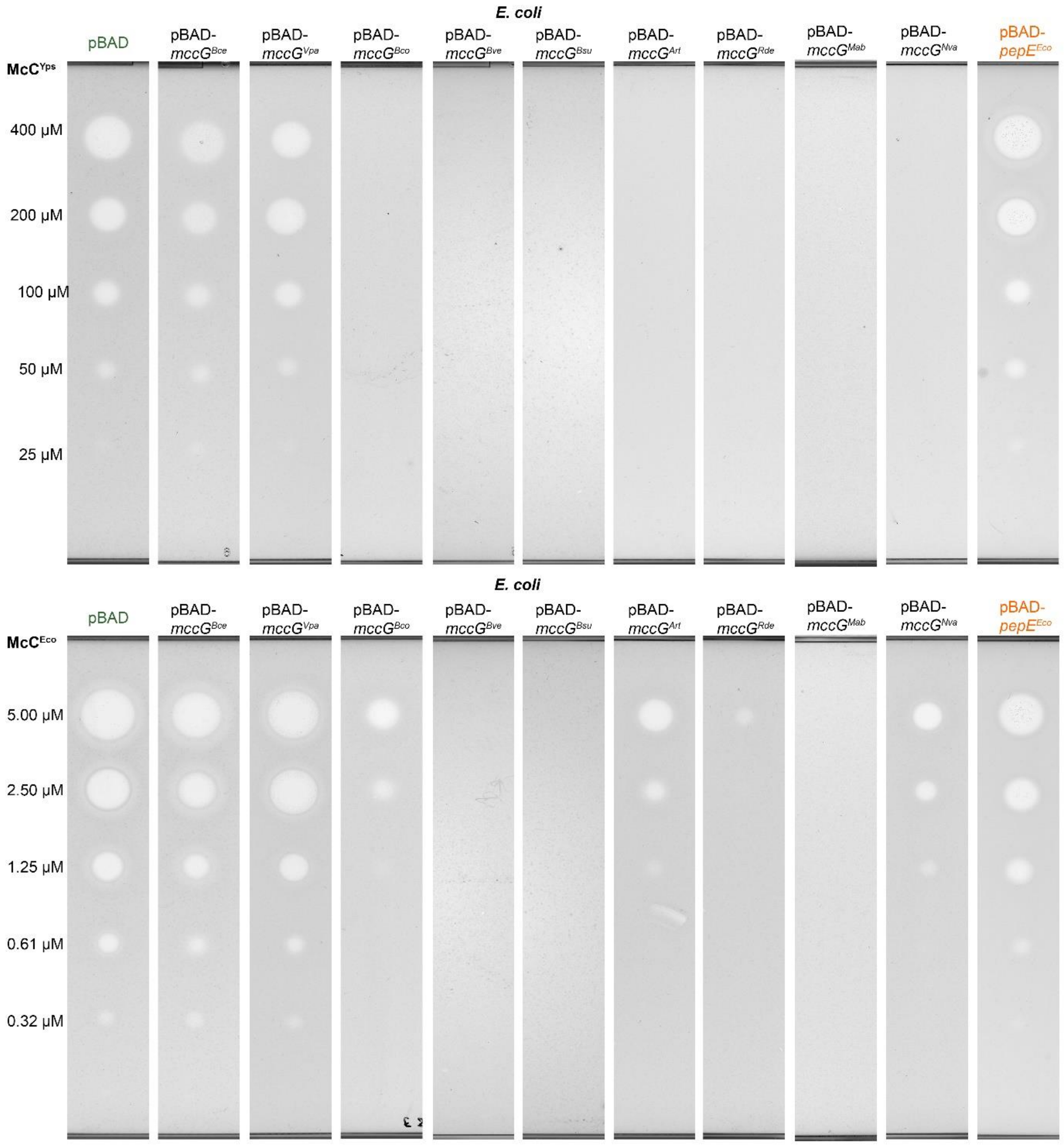


**Figure S3.** Susceptibility to $McC^{Yps}$ and $McC^{Eco}$ of *E. coli* harboring the plasmids with the indicated *mccG*$^{Nva}$ homologs.

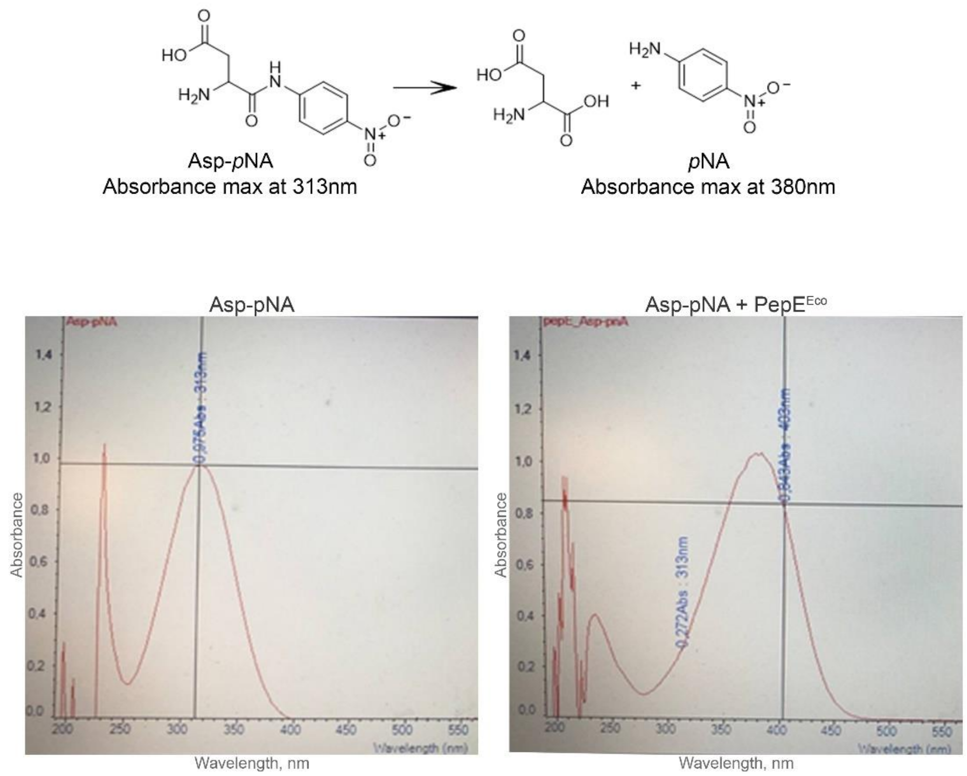


**Figure S4**. Recombinant PepE$^{Eco}$ hydrolyzes synthetic Asp-*p*NA substrate. Absorbance spectra of Asp-pNA (left panel) and Asp-pNA incubated with PepE$^{Eco}$ (right panel).

```
Cph             ------MPLSSQPAILIIGGAEDKVHGREILQTFW----SRSGGNDAIIGIIPSASREPL
PepEse          ------------MELLLLSN--STLPGKAWLEHALPLIANQLNGRRSAV-FIPFAGVTQT
MccGNva         MRRDESAGIDPVRRIILLGGGFSTDPDSLLDEYVL----SASAVDKPRVCFIPTASGDSR
                              ::::..  ..  .    :       .      . : :** *.

Cph             LIG--ERYQTIFSDMGVKELKVLDIRDRAQGDDSGYRLFVEQCTGIFMTGGDQLRLCGLL
PepEse          WDEYTDKTAEVLAPLGVNVTGI----HRVADPLA----AIEKAEIIIVGGGNTFQLLKES
MccGNva         --GYTDRFYSAFTRMNCTPSHLWLFHDRSADMAS----LVANQDILYVGGGSTANLLALW
                     :.    :: :. .   :     *  .  :     : :   : : **.   .*

Cph             ADTPLMDRIRQRVHNGEISLAGTSAGAAVMGHHMIAGGSSGEWPNRALVDMAV--GLGIV
PepEse          RERGLLAPMADRVKRGAL-YIGWSAGANL---ACPTIRTTNDMP---IVDPNGFDALDLF
MccGNva         RLHGLDRLIRDAYRRGVL-LCGISAGAACWFDACLTDSFGDLRP---LKD-----GLGLL
                    *   : :  ..* :   * ****        :    .  *   : *     .*.:.

Cph             PEIVVDQHFHNRNRMARLLSAISTHPELLGLGIDEDTCAMFERDGSVK--VIGQGTVSFV
PepEse          P-LQINPHFTN---------------------ALPEGHKGETREQRIRELLVVAPELTVI
MccGNva         E-GSFCPHFDAEPERDQLYGQAVSDGSLPGGWALQDGAAALFRNEELRDIVTRNGTSTII
                    .  **                          :      *:  :.  :      :.:

Cph             DA-RDMSYTNAA-----LVGANAPLSLHNLRLNILVHGEVYHQVKQRAFPRVT
PepEse          GL-PEGNWIQVSNGQAVLGGPNTTWVFKAGEEAVAL--EAGHRF---------
MccGNva         GMRRENGYTQIS--------------RKFSKAQILT--EPGGES---------
                .   : .: : :               :     :    *
```

**Figure S5.** A multiple sequence alignment of cyanophycinase from *Synechocystis* sp. (WP_010872518.1), the PepE peptidase from *S. enterica* (WP_000421792.1), and MccG$^{Nva}$ from *N. vaccinii* NBRC 15922 (WP_218027309.1) is presented. The alignment was built using MUSCLE with default parameters. Experimentally confirmed catalytic residues of Cph and PepE and are highlighted in red. Based on the sequence alignment, only two of these residues, a serine and a histidine are conserved in all three proteins.

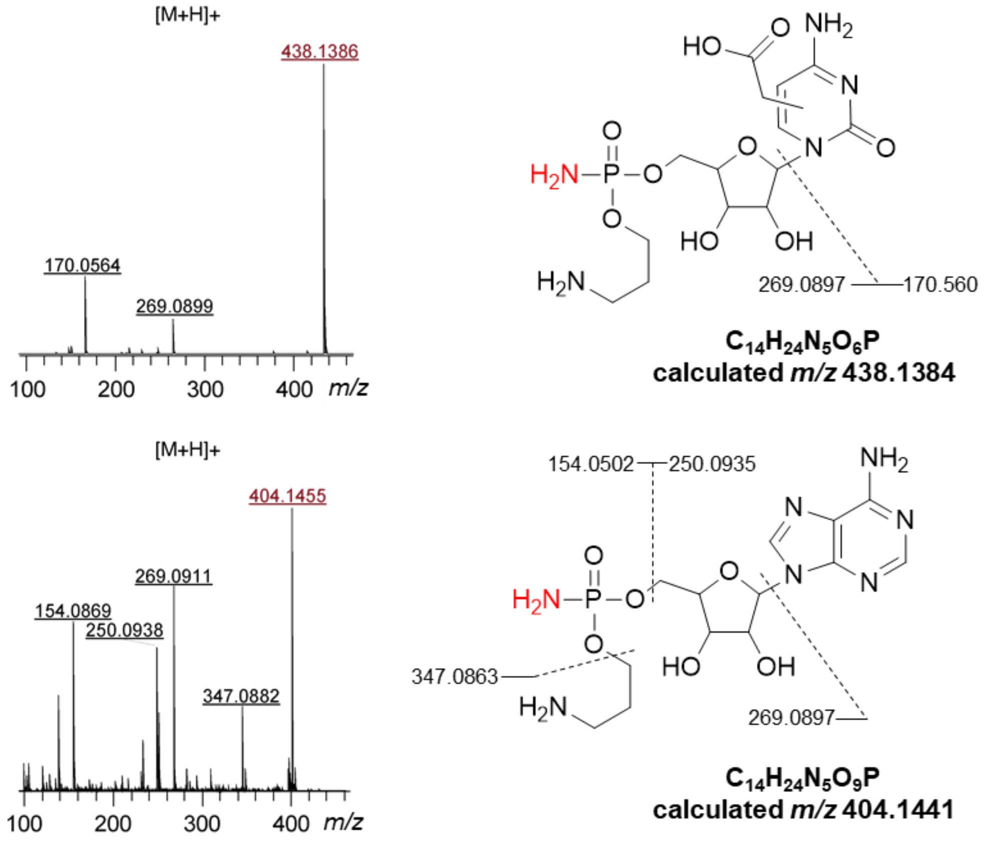


**Figure S6.** ESI-MS/MS fragmentation spectra of the products of MccG$^{Nva}$-mediated hydrolysis of the McC$^{519}$ (upper panels) and McC$^{553}$ (lower panels). Parent ions on mass-spectra are labeled with red color font.

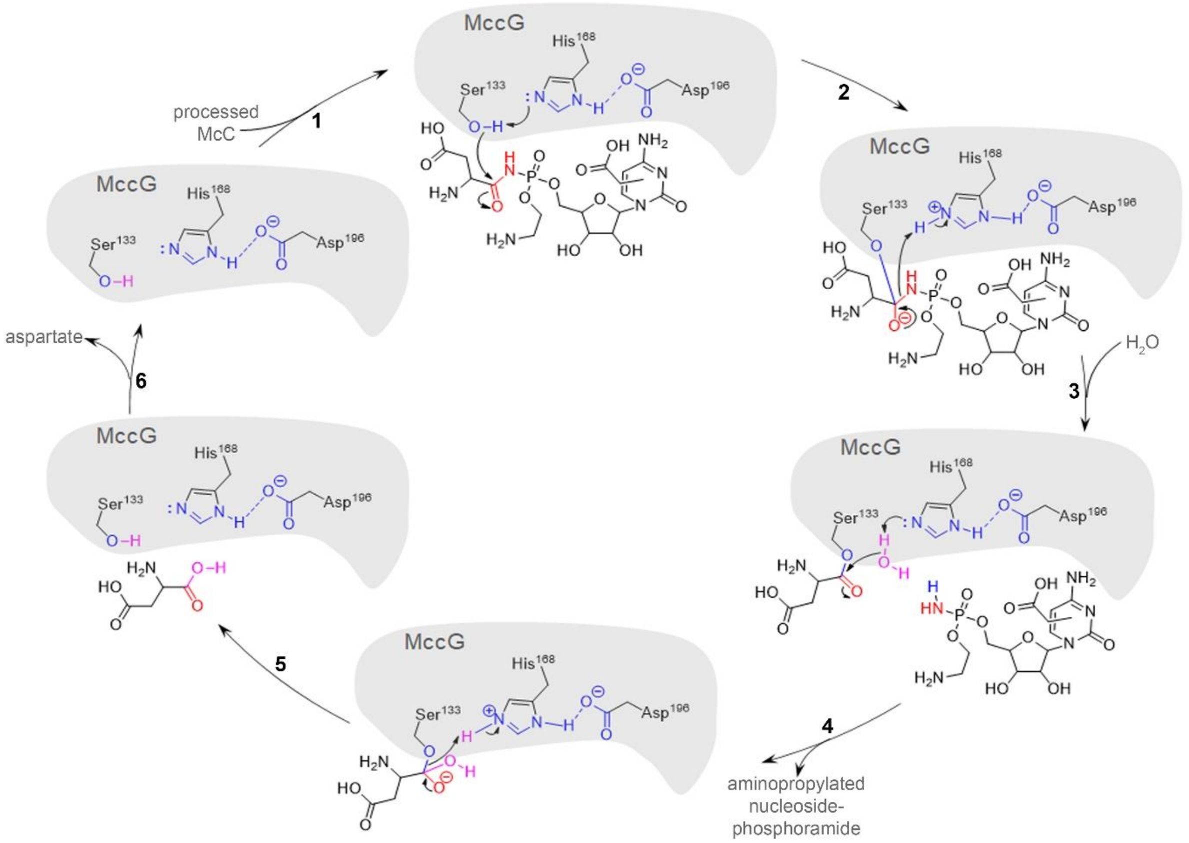


**Figure S7. A proposed reaction mechanism of MccG**. At the first step, Ser133 deprotonated with His168 attacks the carbonyl group of processed McC aspartate forming the first tetrahedral complex. The His168 complex with proton is stabilized via a hydrogen bond with the negatively charged Asp196. The His168 hydrogen is then taken by the nitrogen of the phosphamide bond of the substrate, releasing the nucleotide part. Next, the remaining covalently bound aspartate is attacked by a water molecule, which, similarly to Ser133 at the first step, is deprotonated by His168. The newly formed tetrahedral complex decomposes with the release of free aspartate and the catalytic triad is returned to its initial state.

**Table S1. Primers used in the study**

| No | Primer ID | Sequence 5'-3' | Purpose |
|---|---|---|---|
| 1 | MccB_Nva_F_BamHI | TATTATGGATCCAAACGAATACTTGCAACTG | Cloning into pRSFDuet-1 |
| 2 | MccB_Nva_R_SacI | ATTATAGAGCTCTTAGCGGCCGAGCACGAACGTAAG | |
| 3 | PepE_Eco_F_SalI | ATTTATGTCGACATGGAACTGCTTTTATTGAG | Cloning into pBAD_SalRBS |
| 4 | PepE_Eco_R_HindIII | TATATAAAGCTTAAAAACGGTGACCAGCTTCC | |
| 5 | MccF_F_SalI | TATTATGTCGACATGATACAATCTCATCCACTAC | |
| 6 | MccF_R_HindIII | TATTATAAGCTTATTTCTCGGTAGATAGATATTGTTCTG | |
| 7 | MccG_F_SalI | ATTTATGTCGACATGCGTCGTGACGAGTCGGC | |
| 8 | MccG_R_HindIII | ATATTATAAGCTTATGACTCCCCGCCCGGC | |
| 9 | MccG_Art_F_SalI | ATTTATGTCGACATGGCAGCCCAACAGCCAAC | |
| 10 | MccG_Art_R_HindIII | TATTAAAAGCTTAGCCAAGGAAACGGGCCTCC | |

| | | | |
|---|---|---|---|
| 11 | MccG_Bsu_F_SalI | ATTATTGTCGACATGAAGCAGATTATTGC GATG | |
| 12 | MccG_Bsu_R_HindIII | TATATTAAGCTTACCCTAAATATTTGACC GG | |
| 13 | MccG_Bce_F_SalI | ATTATAGTCGACATGAAATTAGCTGTCAT TGGTG | |
| 14 | MccG_Bce_R_HindIII | TATATAAAGCTTATAAATAGCTTTTTAAA GTAATATC | |
| 15 | MccG_Bco_F_SalI | ATTATTGTCGACATGAGGCAAATTATCGC AATG | |
| 16 | MccG_Bco_R_HindIII | TATATTAAGCTTACATGTCATTATTATCA TCTAAATATTTTA | |
| 17 | MccG_Bve_F_SalI | ATTTATGTCGACATGACATTGAAGCAGAT TATTGC | |
| 18 | MccG_Bve_R_HindIII | ATTTATAAGCTTAAAATTCAGATAACGGT ACGG | |
| 19 | PepE_Eco_F_NdeI | ATTTATCATATGGAACTGCTTTTATTGAG | Cloning into pET22(b) |
| 20 | PepE_Eco_R_XhoI_wsc | TTATTTCTCGAGAAAACGGTGACCAGCTT CCAG | |
| 21 | MccF_F_NdeI | TATTATCATATGATACAATCTCATCCACT AC | |
| 22 | MccF_R_XhoI_wsc | ATTTTATCTCGAGTTTCTCGGTAGATAGA TATTGTTCTG | |

| | | | |
|---|---|---|---|
| 23 | MccG_F_NdeI | TATTTACATATGCGTCGTGACGAGTCGGC | |
| 24 | MccG_R_SalI_wsc | ATATTATGTCGACTGACTCCCCGCCCGGCTC | |
| 25 | MccG_S133A_F | GCTGCTGTGCGGCATCGCCGCCGGTGCCGCCTGCTG | Mutagenesis |
| 26 | MccG_S133A_R | CAGCAGGCGGCACCGGCGGCGATGCCGCACAGCAGC | |
| 27 | MccG_H168A_F | GCAGTTTCTGCCCGGCGTTCGACGCCGAGCCCGAAC | |
| 28 | MccG_H168A_R | GTTCGGGCTCGGCGTCGAACGCCGGGCAGAAACTGC | |
| 29 | MccG_D196A_F | GAGGATGGGCCCTCCAGGCGGGCGCGGCCGCATTGTTTC | |
| 30 | MccG_D196A_R | GAAACAATGCGGCCGCGCCCGCCTGGAGGGCCCATCCTC | |
| 31 | MccG_D196E_F | GAGGATGGGCCCTCCAGGAAGGCGCGGCCGCATTGTTTC | |
| 32 | MccG_D196E_R | GAAACAATGCGGCCGCGCCTTCCTGGAGGGCCCATCCTC | |
